# CircAge: A Comprehensive Resource for Aging-Associated Circular RNAs Across Species and Tissues

**DOI:** 10.1101/2024.11.09.622782

**Authors:** Xin Dong, Zhen Zhou, Yanan Wang, Ayesha Nisar, Shaoyan Pu, Longbao Lv, Yijiang Li, Xuemei Lu, Yonghan He

## Abstract

Circular RNAs (circRNAs) represent a novel class of RNA molecules characterized by a circular structure and enhanced stability. Emerging evidence indicates that circRNAs play pivotal regulatory roles in the aging process. Despite this, there is a lack of a systematic resource that integrates aging-associated circRNA data. Therefore, we developed a comprehensive database named circAge, which encompasses 689 aging-related samples from 7 species and 21 tissue types. We also generated 47 new tissue samples from mice and rhesus monkeys through high-throughput sequencing. Integrating predictions from multiple bioinformatics tools, we identified over 413,378 unique circRNAs. Our data analysis revealed a general increase in circRNA expression levels with age, with approximately 22% of circRNAs demonstrating sequence conservation across species. The circAge database systematically predicts potential interactions between circRNAs and miRNAs, RNA-binding proteins, and assesses the coding potential of circRNAs. This resource lays a foundation for elucidating the regulatory mechanisms of circRNAs in aging. As a comprehensive repository of aging-associated circRNAs, circAge will significantly accelerate research in this field, facilitating the discovery of novel biomarkers and therapeutic targets for aging biology and developing diagnostic and therapeutic strategies for aging and age-related diseases. CircAge is publicly available at https://circage.kiz.ac.cn.

## Introduction

Circular RNAs (circRNAs) are a novel class of RNA molecules characterized by a circular structure, exhibiting higher stability and cell-type-specific expression compared to linear RNAs [1, 2]. Recent studies have demonstrated that circRNAs not only participate in gene expression regulation but also play pivotal roles in crucial biological processes such as cell differentiation, proliferation, and apoptosis [3-6]. With advancements in aging research, accumulating evidence suggests that circRNAs exert critical regulatory functions during the aging process. Research has revealed that circRNAs progressively accumulate in aging tissues, including the heart and brain [7, 8]. circRNAs can modulate aging and age-related diseases through diverse mechanisms. For instance, circHIPK3 acts as a scaffold for HuR and E3 ubiquitin ligase, promoting HuR degradation and consequently influencing cardiac aging [9]. Circ-Foxo3 can bind to aging-associated proteins such as ID-1, E2F1, and FAK, thereby promoting cellular senescence [10]. However, only a limited number of studies have elucidated the mechanisms by which circRNAs contribute to aging. The specific roles and modes of action of most circRNAs in aging and related diseases remain largely unexplored. Therefore, there is an urgent need to establish a comprehensive database of aging-associated circRNAs to facilitate research in this field.

Currently, several public circRNA databases have been established, such as circBase [11], which have cataloged circRNA sequences and annotations from numerous species. Additionally, databases like CSCD2 [12], TSCD [13], and MiOncoCirc [14] have collected circRNA expression and function information from normal and cancer samples, providing valuable resources for investigating the roles of circRNAs in tumorigenesis and other diseases. However, no database has systematically collected aging-associated samples and identified aging-related circRNAs, presenting an inconvenience for research in this field.

Therefore, the establishment of a systematic database for aging-related circRNAs, which integrates expression profiles across diverse tissues and age groups while predicting potential regulatory mechanisms, is an invaluable resource for comprehensively understanding the roles of circRNAs in aging. This database will not only aid in the discovery of novel aging biomarkers and therapeutic targets but also foster interdisciplinary collaborative research. By offering new perspectives for the diagnosis and treatment of aging and age-related diseases, it will significantly advance both basic and clinical translational research on circRNAs within the field of aging biology.

## Methods

### Data sources

We comprehensively collected aging-related tissue and cell line sequencing data from the GEO database. Specifically, we obtained samples from 21 tissue types across 7 species: *Homo sapiens, Macaca mulatta, Mus musculus, Rattus norvegicus, Drosophila melanogaster, Caenorhabditis elegans*, and *Danio rerio*, totaling 689 samples. These datasets include age information or groupings of aging and young samples, and the samples were prepared using total RNA with rRNA depletion, making them suitable for identifying aging-associated circRNAs.

To further expand our database with aging and young tissue samples from mice and rhesus monkeys, we conducted high-throughput sequencing. We collected age-gradient (young, middle-aged, and old) samples of fat, heart, kidney, lung, and spleen tissues from mice, as well as liver and spleen tissues from rhesus monkeys, totaling 47 samples. By accumulating aging data across multiple species and systemically predicting the expression and functions of circRNAs, we aim to comprehensively elucidate the molecular dynamics underlying the aging process (**Figure 1**).

**Figure 1.**
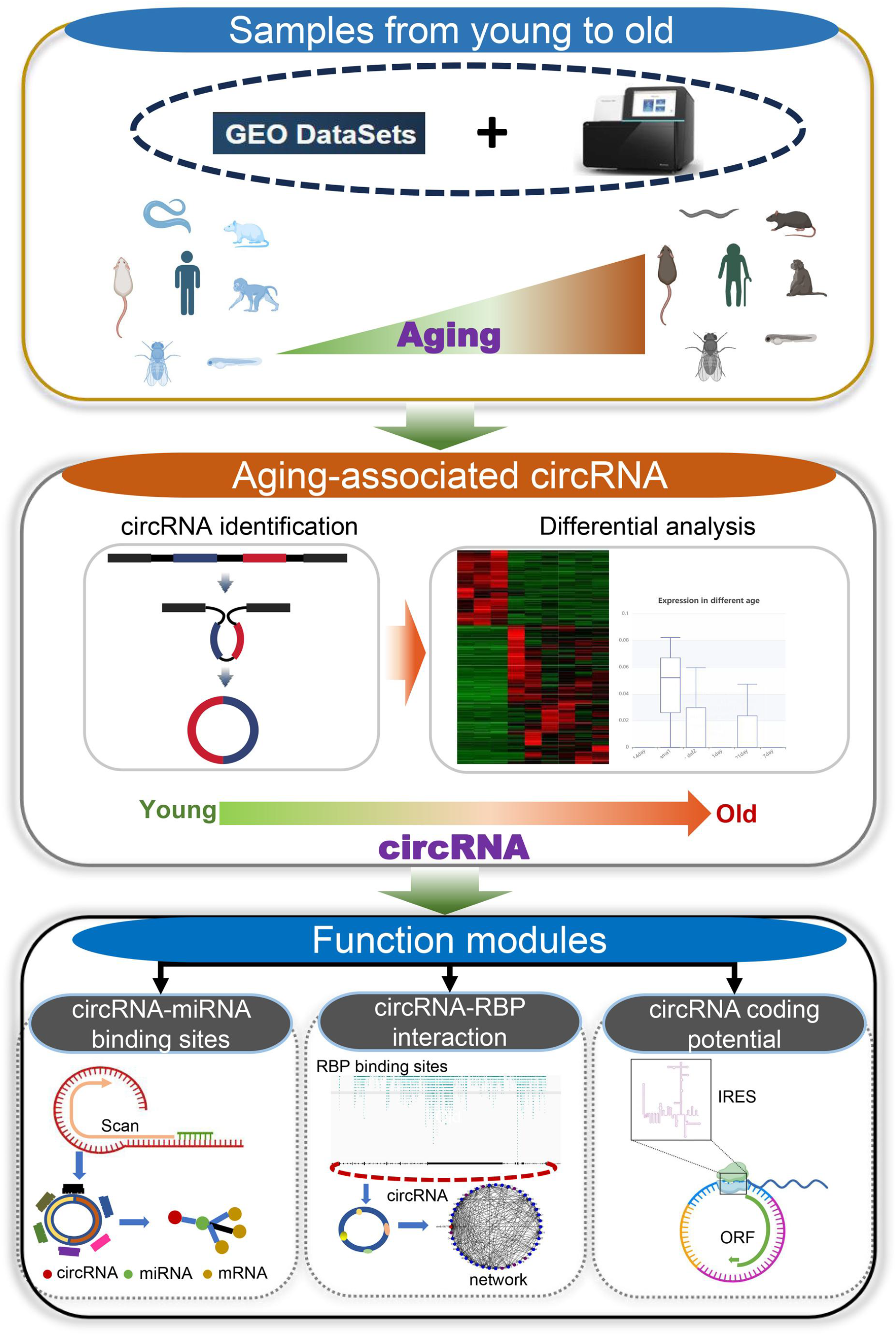
Schematic of the data processing pipeline in the circAge database. The circAge database collects multi-species data from the GEO database and large-scale sequencing in lab. The circAge combines two classic identification tools to detect circRNAs in each sample, and performs differential analysis combined with age information to screen for aging-associated circRNAs. The circAge performs annotation and sequence extraction for all circRNAs by combining gene annotations, and conducts systematic functional predictions such as miRNA/RBP binding sites, circRNA coding potential, circRNA conservation, *etc*.

### Tissue sequencing

Total RNA was extracted from mouse and rhesus monkey tissues, followed by rRNA depletion. The remaining RNA was then reverse-transcribed into cDNA and constructed into strand-specific libraries. High-throughput sequencing was performed on these libraries using the Illumina NovaSeq 6000 system. This study was approved by the Institutional Animal Care and Use Committee of the Kunming Institute of Zoology, Chinese Academy of Sciences, with the approval number IACUC-PA-2023-08-031. All raw sequencing data are available in the Genome Sequence Archive (PRJCA026538) [15].

### circRNA identification

To remove sequencing adapters and low-quality bases, quality control of the raw reads was performed using trim_galore. The clean reads were mapped to the reference genome using the STAR software [16]. CIRCexplorer2 [17] and circRNA_finder [18] were employed for circRNA identification and quantification. We integrated the predictions from both algorithms and normalized the circRNA back-spliced junction (BSJ) counts using the Counts Per Million (CPM) method, calculated as (number of circRNA BSJs * 10^6) / number of mapped reads. Additionally, we annotated the circRNAs with their host genes using in-house scripts and genomic annotation files. Based on the genomic locations, circRNAs can be classified as exonic circRNAs, intronic circRNAs, or intergenic circRNAs. Furthermore, circRNAs were categorized as originating from mRNAs or lncRNAs, depending on whether their parental genes were protein-coding or not.

### Identification of aging-associated circRNAs

For each tissue, we combined the normalized circRNA expression levels into a matrix. Utilizing the grouping information, we performed differential expression analysis with t-tests. circRNAs exhibiting significant differential expression (*p-*value <= 0.05) between age groups were identified as aging-associated circRNAs.

### circRNA conservation analysis

The circAge database encompasses circRNA data from multiple species. To assess the conservation of these circRNAs, we first reconstructed their full-length sequences using previously established methods [11]. Then we extracted a 100 bp window (±50 bp) surrounding the back-splice site of each circRNA, and we performed BLAST searches to compare the junction sequences across tissues and species [19]. Subsequently, we filtered the alignment results, retaining those with an E-value less than or equal to 10^-5, identity greater than 80% and alignment length greater than 80. circRNAs with matches in multiple species were considered conserved.

### circRNA-miRNA binding prediction

To provide potential miRNA binding sites within the circRNA sequences, we scanned the full-length circRNA sequences using the miRanda and TargetScan algorithms to obtain all potential miRNA binding locations [20, 21]. Additionally, to facilitate downstream target prediction for users, we integrated validated miRNA-target gene data from the Tarbase database [22], allowing users to conveniently query circRNA-miRNA-target gene interaction networks. Furthermore, to enable users to explore the downstream functions of miRNAs, we performed Gene Ontology (GO) enrichment analysis on all target genes with the cluster profiler package [23].

### circRNA RBP binding site prediction

We obtained CLIP-seq data for 4 species (*H. sapiens, M. musculus, D. melanogaster, C. elegans*) from the POSTAR3 database [24]. Using this data, we predicted potential RNA-binding proteins (RBPs) that may interact with circRNAs based on previous methods [25], retaining RBP binding sites within the circRNA sequences or within 1 kb upstream or downstream regions.

### circRNA coding potential prediction

circRNAs have been shown to encode functional peptides. In circAge, we comprehensively predicted the coding potential of all circRNAs. We employed the CPAT algorithm to assess the coding potential of circRNA sequences and further used IRESfinder to detect the presence of internal ribosome entry sites (IRES) within the circRNA sequences [26-28].

### Database construction

The circAge database was deployed in a virtual machine running Windows server 2016 using Internet Information Services (IIS) for service. The backend of the circAge database was developed using C# language and NET MVC framework, meanwhile the web frontend was organized using HyperText Markup Language (HTML), Cascading Style Sheets (CSS), JavaScript and LayUI framework. All those data were stored in Microsoft SQL Server. As massive amount of data with over 200 million entries, Apache Solr facilitated its blazing-fast search. MinIO, a S3-compatible object storage system, provided associated file for downloading. The interactive visualization of the analysis results was implemented using jQuery, Apache ECharts (https://echarts.apache.org/) and D3.js (https://d3js.org/).

## Results

### A comprehensive database of aging-related circRNAs across 7 species and 21 tissues

We collected a total of 689 aging-related samples from multiple GEO datasets, encompassing 7 species (*H. sapiens, M. mulatta, M. musculus, R. norvegicus, D. melanogaster, C. elegans*, and *D. rerio*) and 21 tissue types (dataset list is shown in Supplementary Table S1). Additionally, we generated 47 new RNA-seq samples, including tissues (fat, heart, kidney, lung, and spleen) from young, middle-aged, and old mice, as well as rhesus monkey liver and spleen samples. After quality control, we obtained 2.7 billion raw reads, with an average of 30 M reads per sample (**Figure 2A**). By integrating the circRNA predictions from CIRCexplorer2 and circRNA_finder, we identified a total of 413,378 unique circRNAs across all samples. The identified circular RNAs originated from different genomic regions, with 69.52% being exonic circRNAs, 23.55% intronic circRNAs, and 6.94% intergenic circRNAs (**Figure 2B**). Both protein-coding and non-coding genes contributed to circRNA biogenesis, with 96% circRNAs derived from mRNAs and 3% from lncRNAs (**Figure 2C**).

**Figure 2.**
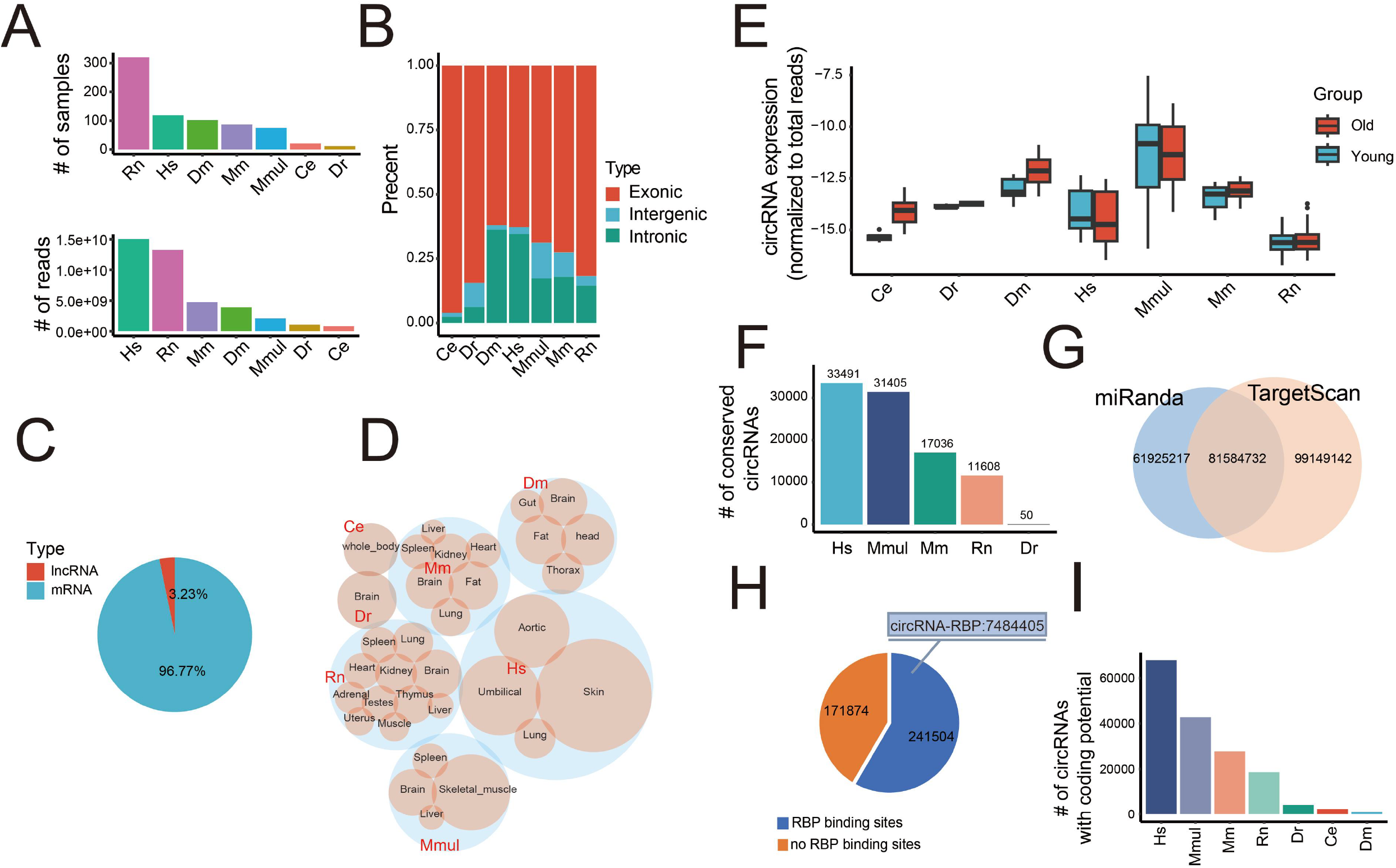
Data statistics and summary of circAge database. **A**. Number of collected samples and sequencing reads in each species. Hs: *Homo sapiens*; Mm: *Mus musculus*; Mmul: *Macaca mulatta*; Rn: *Rattus norvegicus*; Dm: *Drosophila melanogaster*; Dr: *Danio rerio*; Ce: *Caenorhabditis elegans*. **B**. Proportion of circRNAs in different genomic locations. **C**. Proportion of circRNAs in different host gene type. **D**. Number of circRNAs detected in various species and tissues, with larger bubbles representing higher abundance. **E**. Overall circRNA expression (normalized to total reads) profiles across different species. **F**. Number of conserved circRNA in different species. **G**. Number of predicted miRNA binding sites in circRNA with miRanda and TargetScan. **H**. Number of circRNAs harboring RBP binding sites, with the upper annotation indicating the total number of circRNA-RBP interactions. **I**. Number of circRNAs with coding potential in different species.

The number of detected circRNAs varied across species, tissues (**Figure 2D**). Overall, human samples yielded more circRNAs compared to other species. Brain tended to exhibit higher circRNA abundances. We performed differential expression analysis for circRNAs in each tissue to identify aging-associated circRNAs. In total, 17527 circRNAs were found to be significantly associated with aging (*p* ≤ 0.05) across the 21 different tissues. Interestingly, in multiple tissues, circRNA expression showed an increasing trend during the aging process (**Figure 2E**).

### circRNA conservation and functional annotation

We performed sequence alignments to identify conserved circRNAs across different species, considering circRNAs with similar sequences as conserved. The results revealed 93,590 conserved circRNAs, accounting for an average of 22% of the total circRNAs in each species (**Figure 2F**). This finding suggests a high degree of species-specificity for circRNAs, consistent with previous studies [25].

Furthermore, we obtained mature miRNA sequences and IDs from the Targetscan website and used the Targetscan and miRanda algorithms to predict miRNA binding sites within the full-length circRNA sequences. Our analysis identified a substantial number of 242,659,091 circRNA-miRNA binding sites (**Figure 2G**). Additionally, we integrated the validated miRNA target genes from Tarbase database and performed GO enrichment analysis on the downstream target genes of miRNAs, facilitating user research. For circRNA-RBP interactions, we integrated data from the POSTAR3 database and obtained numerous circRNA-RBP interaction sites. The results indicate that most circRNAs harbor RBP binding sites (**Figure 2H**). Meanwhile, the circAge database has predicted the coding potential of circRNAs, and the results show that a large number of circRNAs have coding potential (**Figure 2I**).

### Database Access

circAge provides a user-friendly web interface. The Home page mainly shows the introduction to the circAge database, basic statistics, and data summary. Clicking on a specific species will direct the user to the circRNA search interface for that species. Users can also perform quick searches for circRNAs, miRNAs, and RBPs on the Home page (**Figure 3**).

**Figure 3.**
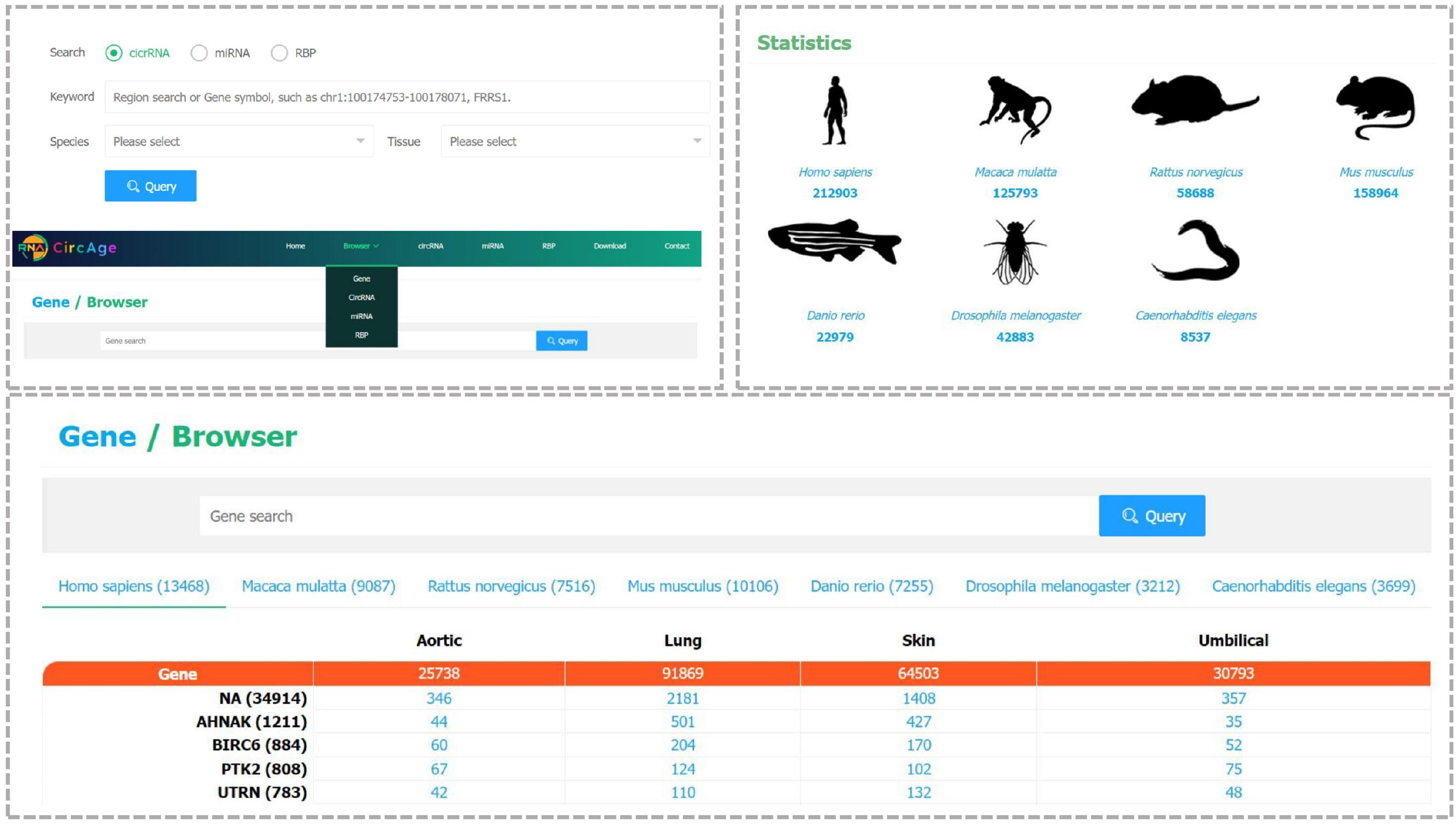
Basic search function of the circAge database homepage and browser pages.

On the Browser page, users first need to select whether to browse genes, circRNAs, miRNAs, or RBPs module. The page will then generate a matrix by species, with rows representing genes, circRNAs, miRNAs, or RBPs, and columns representing tissues. The numbers in the matrix indicate the number of records for that gene, circRNA, miRNA, or RBP in circAge. Clicking on rows names will navigate to the search results for that entry, while clicking on a number in the matrix will redirect to the search page for that gene, circRNA, miRNA, or RBP in the corresponding tissue (**Figure 3**).

The circAge database consists three modules: circRNAs, miRNAs, and RBPs. In circRNA panel, users can search for circRNAs by coordinates or browse by selecting a specific species and tissue type. The table below displays detailed information about each circRNA, including its ID, species, host gene, and circRNA type. Users can click on the species, tissue, and host gene columns to perform further searches for the corresponding terms. Additionally, clicking on a circRNA entry will redirect to its detail page, which displays the circRNA’s main information. The top section shows a schematic diagram of the circRNA, with the upper panel illustrating the structure of the host gene and the circRNA generation site, and the lower left panel display the circRNA structure and potential miRNA binding sites. The lower right part shows the circRNA expression levels across different age groups. Below, the page presents predicted circRNA-miRNA binding results, circRNA-RBP binding predictions, and circRNA coding potential predictions (**Figure 4**).

**Figure 4.**
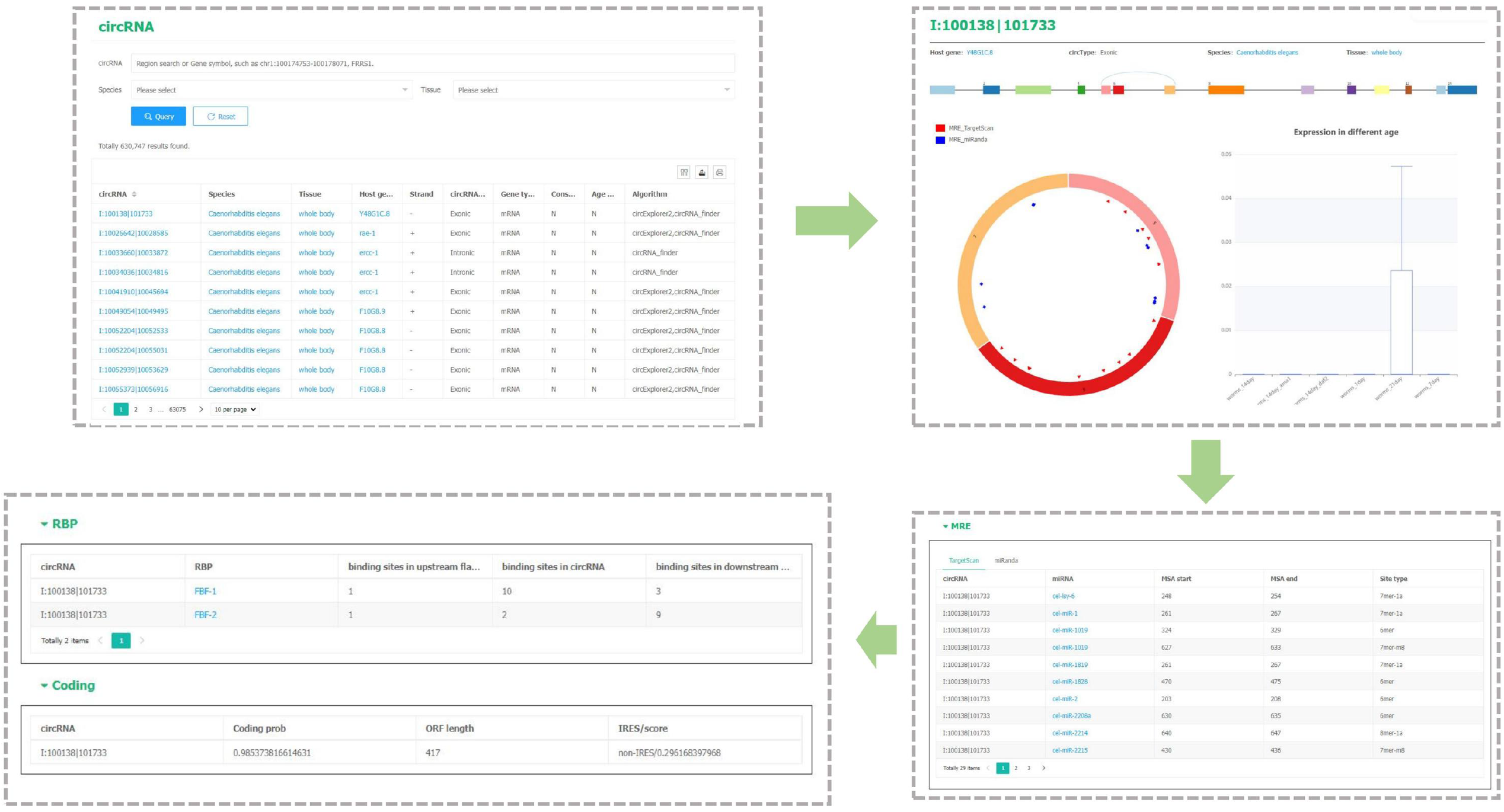
The circRNA detail page of the circAge database displays circRNA search, circRNA annotation, circRNA structure, circRNA expression, miRNA and RBP binding sites within circRNAs, as well as circRNA coding potential prediction results.

On the miRNA page, users can search for specific miRNAs of interest. circAge will return all circRNAs that interact with the queried miRNA. Clicking on a circRNA will display its detail page, while clicking on a species or tissue will navigate to the search interface. Clicking on a miRNA will return a list of its validated target genes and Gene GO enrichment results, allowing users to explore the circRNA-miRNA-mRNA regulatory network and potential downstream functions (**Figure 5**).

**Figure 5.**
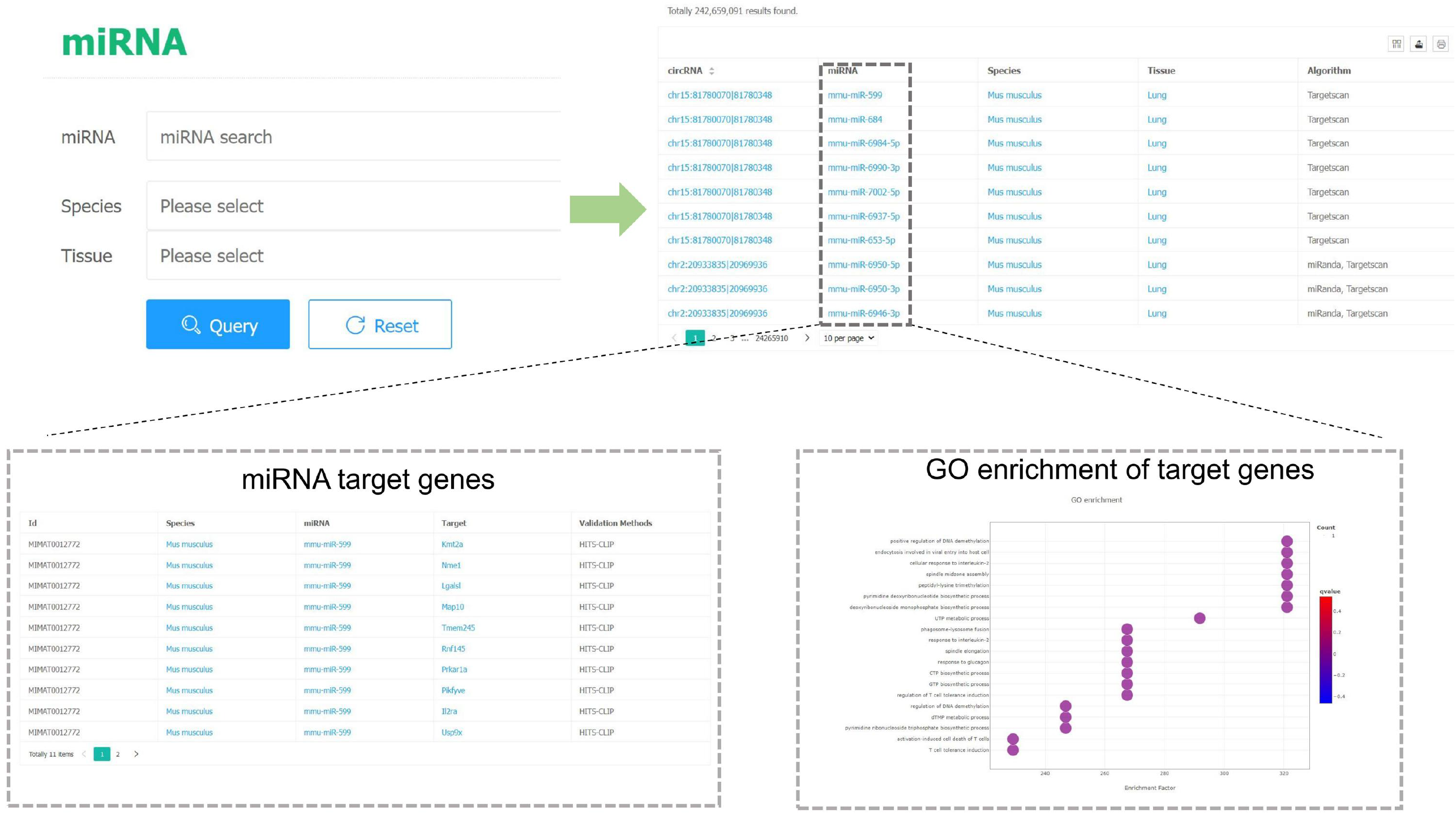
The miRNA module in the circAge database provides a miRNA search function, while integrating downstream target genes of miRNAs and functional analysis results of the target genes.

On the RBP page, users can search for specific RBPs, and circAge will return all circRNAs that bind to the queried RBP. Users can also click on the RBP column, and circAge will display the structure of the corresponding host gene and the circRNA generation site, as well as all binding sites for the RBP within that gene region. All circRNA information in the circAge database is openly accessible, and users can download data through the Download page (**Figure 6**).

**Figure 6.**
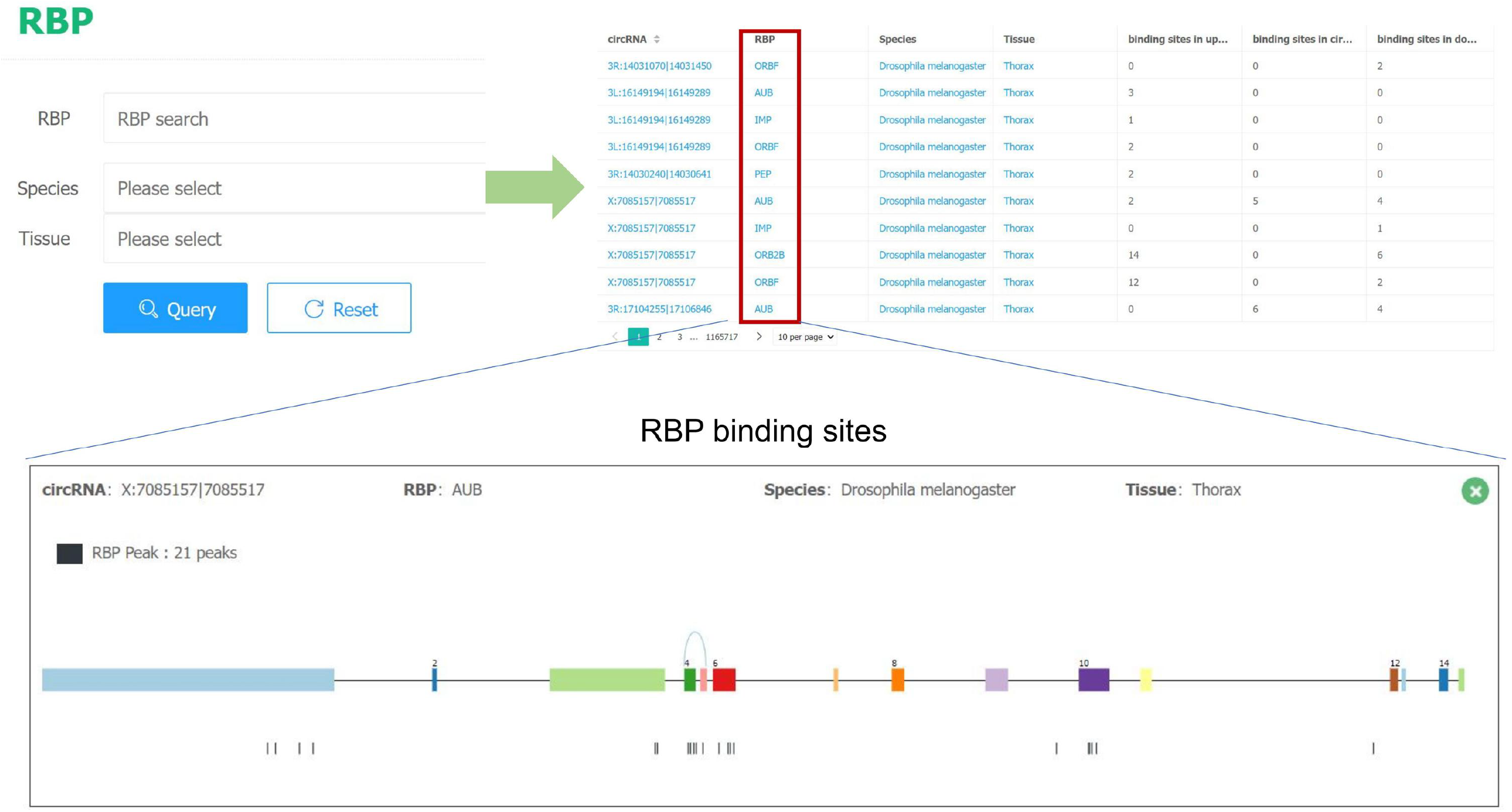
The RBP module in the circAge database provides an RBP search function, while allowing users to view the specific binding sites of RBPs on circRNAs and genes.

### Case study: circCCNB1 functions as a miRNA sponge to regulate aging

To investigate the regulatory roles of circRNAs in aging processes of lung tissue, we queried the CircAge database. Interestingly, circCCNB1 (chr5:69174876|69175537) was found to be significantly downregulated during aging (**Figure 7A**), suggesting its potential regulatory function. Further analysis revealed that circCCNB1 is derived from exons 6 and 7 of the CCNB1 gene and harbors numerous miRNA binding sites within its sequence. By integrating the circRNA-miRNA and miRNA-target gene interaction data from CircAge, we reconstructed a comprehensive regulatory network for circCCNB1 (**Figure 7B-C**).

**Figure 7.**
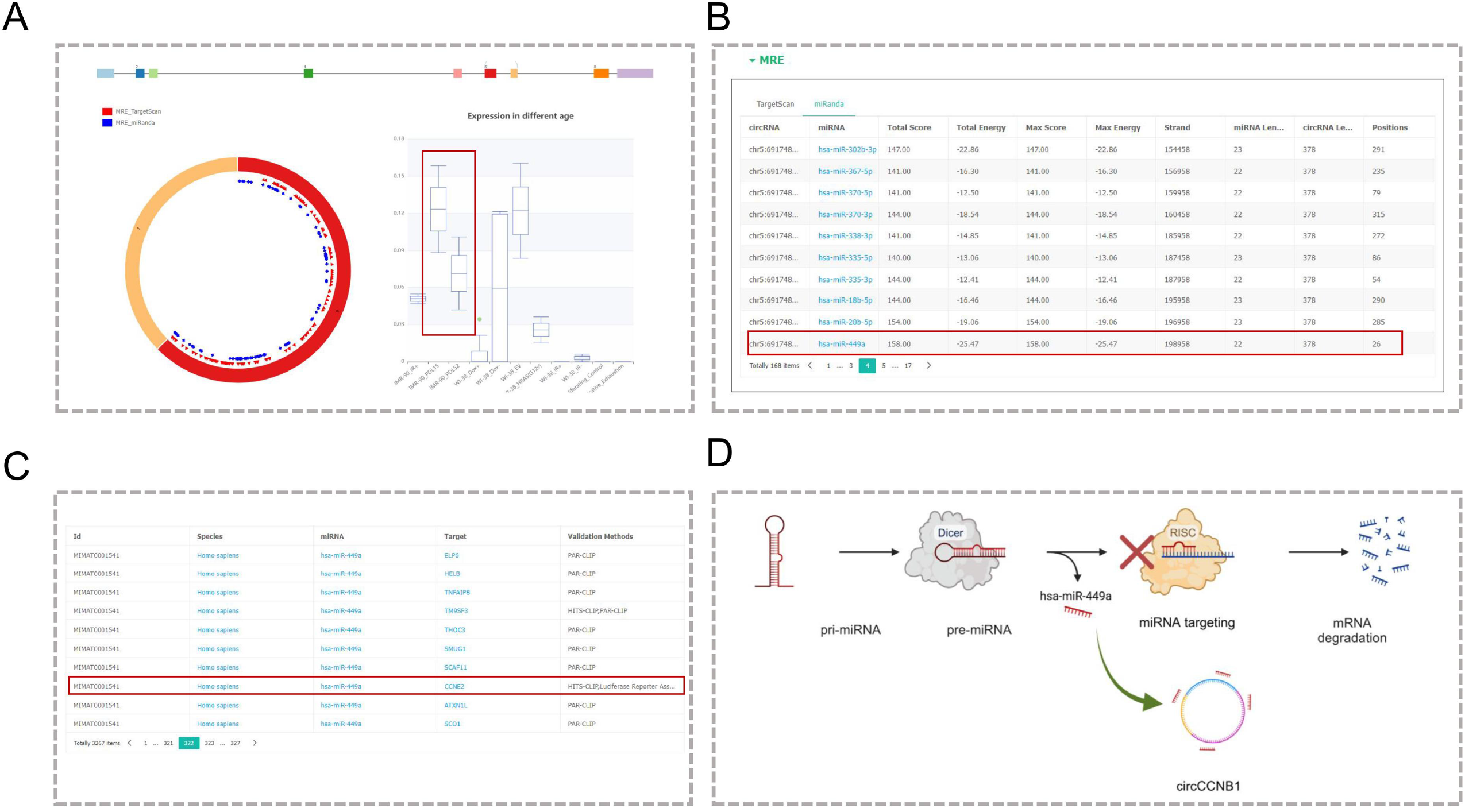
circCCNB1 function prediction using circAge. **A**. circCCNB1 was down-regulated in aging lung tissue. **B**. circCCNB1 harbors multiple miRNA binding sites including has-miR-449a. **C:** has-miR-449a has numerous experimentally validated downstream target genes, such as CCNE2. **D**. The circCCNB1-has-miR-449-CCNE2 axis has been validated to play a role in the regulation of aging.

Literature survey unveiled that the anti-aging role of circCCNB1 has been experimentally validated. It can act as a molecular sponge for hsa-miR-449a, alleviating the repressive effect of this miRNA on its target CCNE2, thereby inhibiting cellular senescence (**Figure 7D**) [29].

## Discussion

This study established a comprehensive database of aging-associated circRNAs, providing a critical resource for investigating the regulatory roles of circRNAs during the aging process. The circAge database encompasses data from multiple species and tissue types, systematically integrating circRNA expression profiles, conservation, potential interactions with miRNAs/RBPs, and coding potential, laying the foundation for elucidating the functions of circRNAs in aging biology. Within the database, circRNA expression levels exhibit an overall upward trend with increasing age, suggesting that circRNAs may be broadly involved in regulating the aging process. Approximately 22% of circRNAs exhibit sequence conservation across different species, and these conserved circRNAs warrant focused attention and in-depth investigation, aiding in the elucidation of conserved regulatory mechanisms underlying the aging process.

The circAge database continuously integrates the latest research developments, creating an extensive circRNA knowledge base that facilitates the elucidation of the complex relationship between circRNAs and aging. The inception of circAge holds substantial significance, expediting research into the regulatory mechanisms of circRNAs within aging biology and serving as a reference for investigating circRNA functions in other biological processes. In the future, combined with the incorporation of experimental validation and complementary studies, the pivotal roles of circRNAs in aging and associated diseases will be further revealed. This will establish a foundation for the development of advanced diagnostic and therapeutic strategies.

## Data availability

CircAge is freely available at https://circage.kiz.ac.cn/.

## Acknowledgments

This work was supported by grants from the National Key R&D Program of China (2023YFC3603300 to Yonghan He), Postdoctoral Fellowship Program of China Postdoctoral Science Foundation (CPSF) (GZC20232763 to Xin Dong), Yunnan Technology Innovation Talent Program (202305AD160021 to Yanan Wang), State Key Laboratory for Conservation and Utilization of Bio-Resources in Yunnan (2023KF008 to Xin Dong), National Natural Science Foundation of China (82171558 to Yonghan He), Yunnan Fundamental Research Projects (202305AH340006 and 202201AS070038 to Yonghan He, 202401CF070064 to Xin Dong), and Cyber Security and Informatization Special Project of Chinese Academy of Sciences (CAS-WX2022SDC-SJ02 to Xuemei Lu). Yonghan He is supported by the Pioneer Hundred Talents Program of the Chinese Academy of Sciences and the Yunnan Revitalization Talent Support Program Young Talent Project. Xuemei Lu is supported by the Yunnan Revitalization Talent Support Program Yunling Scholar Project. Yanan Wang is supported by Technology Support Talent Program of Chinese Academy of Sciences.

